# Evaluation of heat inactivation of human norovirus in freshwater clams using human intestinal enteroids

**DOI:** 10.1101/2022.01.27.477812

**Authors:** Tsuyoshi Hayashi, Yoko Yamaoka, Atsushi Ito, Takashi Kamaishi, Mary K. Estes, Masamichi Muramatsu, Kosuke Murakami

## Abstract

Foodborne disease attributed to consumption of shellfish contaminated with human norovirus (HuNoV) is one of many global health concerns. Our study aimed to determine conditions of heat-inactivation of HuNoV in freshwater clams (Corbicula japonica) using a recently developed HuNoV cultivation system employing stem-cell derived human intestinal enteroids (HIEs). We first measured the internal temperature of clam tissue in a water bath during boiling at 90 °C and found that approximately 2 minutes are required for the tissue to reach 90 °C. Next, GII.4 HuNoV was spiked into the center of the clam tissue followed by boiling at 90 °C for 1, 2, 3, or 4 minutes. The infectivity of the HuNoV in clam tissue homogenates was evaluated using HIEs. We demonstrated that HuNoV in unboiled clam tissue homogenates replicated in HIEs, whereas infectivity was lost in all boiled samples, indicating that heat treatment at 90 °C for 1 minute is sufficient to inactivate HuNoV in freshwater clams. To our knowledge, this is the first study to determine the infectivity of HuNoV tolerability in shellfish using HIEs and our results will be informative to develop strategies to inactivate HuNoV in foods.

## Introduction

Food borne diseases attributed to consumption of unsafe foods, which are contaminated with pathogens (bacteria, viruses, or parasites) or toxic chemical substances, cause 600 million people illness worldwide every year and therefore pose a major public health concern (WHO, 2020; Lee and Yoon, 2021). Foodborne disease is caused by several pathogens including Campylobacter, Salmonella, Listeria, hepatitis A virus and human norovirus (HuNoV). Among those, HuNoV is a most frequently detected pathogen in contaminated foods being consumed by ill individuals (WHO, 2015; Lee and Yoon, 2021). Shellfish, especially oysters, are recognized as one of the major sources for HuNoV-associated foodborne disease due to the accumulation of HuNoVs in the digestive gland by filter feeding (Ludwig-Begall et al., 2021).

The most common inactivation method of pathogens in contaminated foods is to cook them properly at high temperature. Given the varied thermal tolerability of each pathogen, establishment of tailor-made strategies to inactivate respective pathogens is necessary to reduce the risk of foodborne illness. Concerning HuNoV, there was no robust HuNoV cultivation system developed until recently. Therefore, investigations on food inactivation of HuNoV have been carried out by measuring copy numbers of HuNoV genome or infectious virus titers of surrogate viruses such as murine norovirus (MNV) and feline calicivirus (FCV) (Sow et al., 2011; Bozkurt et al., 2015). However, it remains unclear whether these indirect measures reflect inactivation of HuNoV infectivity.

Recently, several HuNoV cultivation systems have been developed, including B cells (Jones et al., 2014), tissue stem cell-derived human intestinal enteroids (HIEs) (Ettayebi et al., 2016; Ettayebi et al., 2021), human induced pluripotent stem cell-derived intestinal epithelial cells (iPSC-derived IECs) (Sato et al., 2019) and zebrafish (Van Dycke et al., 2019). Studies using our HIE system as well as human iPSC-derived IECs showed HuNoV inactivation by heating or disinfectants (alcohol or chlorine) (Ettayebi et al., 2016; Costantini et al., 2018; Sato et al., 2020). However, to our knowledge, an investigation on HuNoV’s inactivation in foods such as bivalves using the HuNoV cultivation system has not been performed so far.

In this study, we evaluated thermal inactivation conditions of freshwater clams artificially contaminated with HuNoV by measuring infectious HuNoV using the HIE culture system.

## Materials and methods

### Measurement of temperature kinetics in freshwater clams subjected to heat treatment

Live freshwater clams (Corbicula japonica) were purchased at a grocery store and maintained overnight at room temperature in diluted seawater. A whole clam body was then taken from its shell and transferred into a 1.5 mL tube. The weights of the clam bodies ranged from 0.16 g to 0.41 g (mean ± standard deviation [SD], 0.29 ± 0.07). To monitor the internal and external temperature of the samples, 2 thermometers were used; a probe thermometer was inserted into a clam body (Fig. 1A) and another was immersed in a water bath set at 90 °C (Fig. 1B). Both temperatures were recorded every 15 seconds up to 5 minutes.

**Figure 1.**
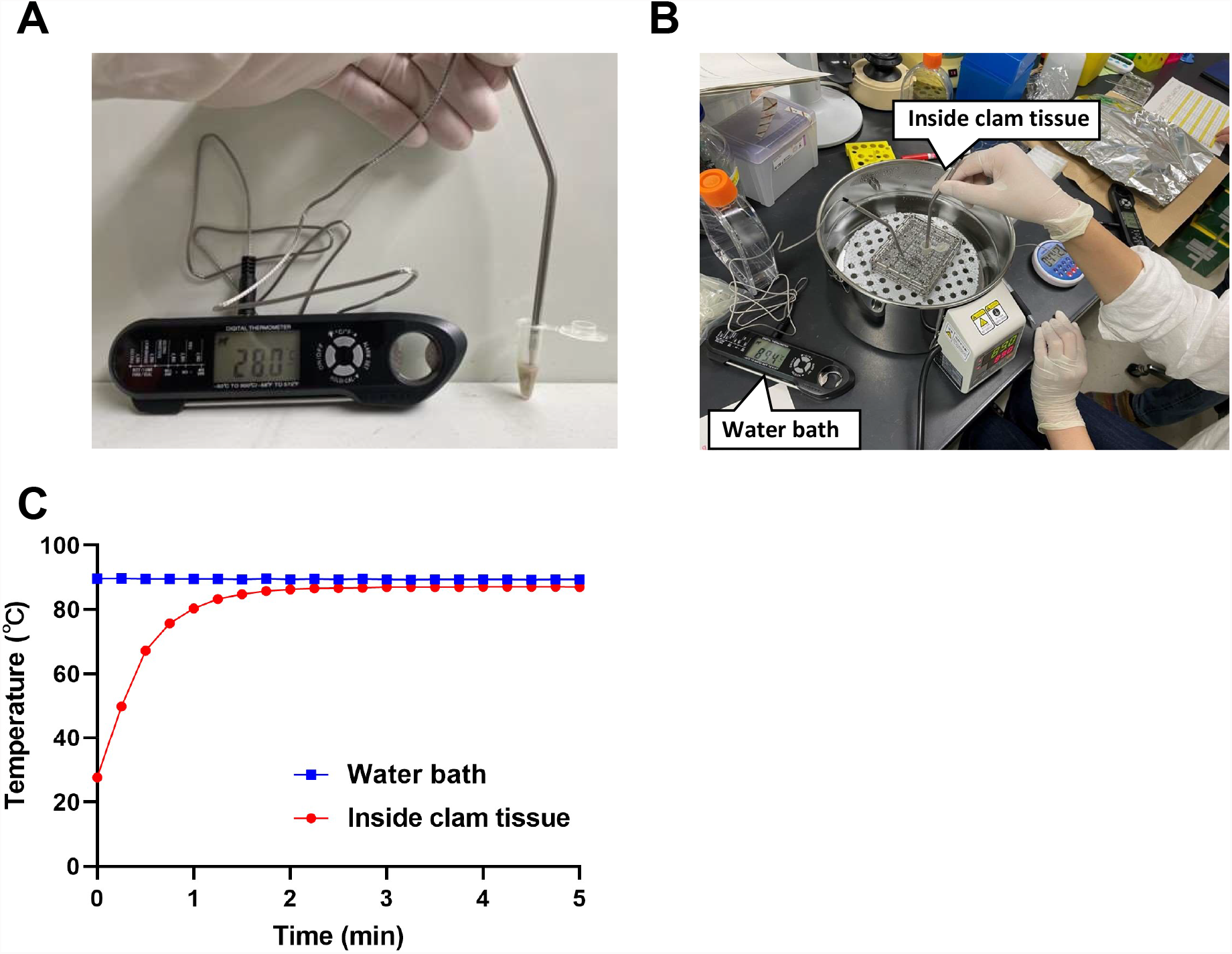
Kinetics of internal temperature in clams subjected to heat treatment. A probe of thermometer was inserted into a clam body in a 1.5 ml tube to measure the internal temperature of the clam (A). A second thermometer probe was put in the water bath to measure the temperature outside the 1.5 ml tube (B). The internal clam tissue temperature and the external water bath temperature were recorded every 15 seconds up to 5 minutes (C). Results were shown as mean ± standard deviation calculated from 5 independent experiments (n=5).

### Artificial inoculation of HuNoV into freshwater clams followed by heat treatment and sample processing

Six freshwater clams were used for each experiment. The clam bodies taken from their shell were injected with 30 μL of either PBS as a control or 10% stool filtrate containing 1.06 × 10^8^ genome equivalents (GEs) of GII.4 [GII.P16] HuNoV (Hayashi et al., 2021) using a 50 μL microsyringe with a fine needle. The artificial inoculated samples were then left-untreated or heat-treated at 90 °C for 1, 2, 3, or 4 minutes, as described above. After cooling the samples down to room temperature, 170 μL of chilled complete medium without growth factors [CMGF(-)] used for culturing HIEs was added to the each sample. The clam bodies were then chopped with scissors and homogenized using a hand mixer, followed by centrifugation at 9,100 x g for 3 min at 4 °C. The supernatant was collected and repeated the centrifugation with the same condition to remove debris. The collected supernatant was stored at -80 °C until used for determining virus infectivity and viral recovery efficiency.

### Evaluation of infectivity of HuNoV in clam extracts using HIEs

HIE culture and HuNoV infection were performed as described previously (Ettayebi et al., 2016; Murakami et al., 2020; Hayashi et al., 2021). Briefly, a jejunal HIE (J2) culture, provided from Baylor College of Medicine under the Material Transfer Agreement, was maintained and propagated as matrigel-embedded, 3-dimensional (3D)-HIEs in complete medium with growth factors [CMGF(+)] or IntestiCult Organoid Growth Medium (Human, STEMCELL).

To prepare monolayer HIE cultures for HuNoV infection, the 3D HIEs were dissociated with TrypLE Express (Thermo Fisher), and seeded onto collagen IV-coated 96-well plates at the number of approximately ∼10^5^ cells/well in the CMGF(+) or IntestiCult media supplemented with ROCK inhibitor Y-27632 (10 μM, Sigma) for 2 days.

The medium was then replaced with IntestiCult Organoid Differentiation Medium (Human, STEMCELL) and the cultures were maintained for additional 2 days. The monolayer HIEs were then inoculated with the clam samples diluted 1:20 in a final volume of 100 μL of CMGF(-) medium in the presence of 500 μM GCDCA, which promotes GII.4 HuNoV infection (Ettayebi et al., 2016). After 1 h incubation at 37 °C, the cells were washed twice with CMGF(-) and further incubated in IntestiCult Organoid Differentiation Medium containing 500 μM GCDCA until 24 hours post infection (hpi). The cells and medium were then collected and subjected to RNA extraction using the Direct-zol RNA MiniPrep kit (Zymo Research) following the manufacturer’s instructions. HuNoV RNA genome equivalents (GEs) were determined by reverse transcription-quantitative PCR (RT-qPCR) analysis using TaqMan Fast Virus 1-Step Master Mix (Thermo Fisher) and GII specific primer/probe sets (Kageyama et al., 2003).

### Evaluation of viral recovery efficiency after homogenization

The RNA was extracted from 5 μL of the inoculum containing 30 μL of HuNoV-containing stool filtrates and 170 μL of CMGF(-) or the HuNoV-contaminated clam extracts, and subjected to RT-qPCR analysis to measure HuNoV GEs. The percentage of recovery efficiency was calculated as HuNoV GEs in clam samples relative to that in the inoculum.

### Statistical analysis

Statistical analysis was performed with ANOVA followed by Dunnett’s multiple-comparison test or two-tailed Student *t* test using GraphPad Prism 9 software. *P* values of < 0.05 was considered statistically significant.

## Results

Heat treatment experiments were carried out using a water bath at 90 °C. Measurements of the internal and external temperatures during heating showed that while the water temperature was stable at around 90 °C during the heating process, the temperature of the clam body required 2 minutes before reaching a stable temperature at 90 °C (Fig. 1C).

We then evaluated the effect of different times of heat inactivation on the HuNoV-contaminated clams. The clam bodies were either spiked with PBS as a non-infection control or HuNoV using microsyringe. The inoculated clam tissue, either left-unheated or heated at 90 °C for 1, 2, 3, or 4 minutes was homogenized. We first evaluated recovery efficiency of spiked HuNoV in clam homogenates (Fig. 2A). Approximately 60% of the inoculated HuNoV GEs (62.0 ± 7.3%) were recovered from HuNoV-spiked clam homogenates without heating, whereas the treatment at 90 °C for 1 minute significantly reduced the recovery efficiency (27.7 ± 15.9%) as compared to the unheated samples. The longer heat treatment further reduced the efficiency (Fig. 2A).

**Figure 2.**
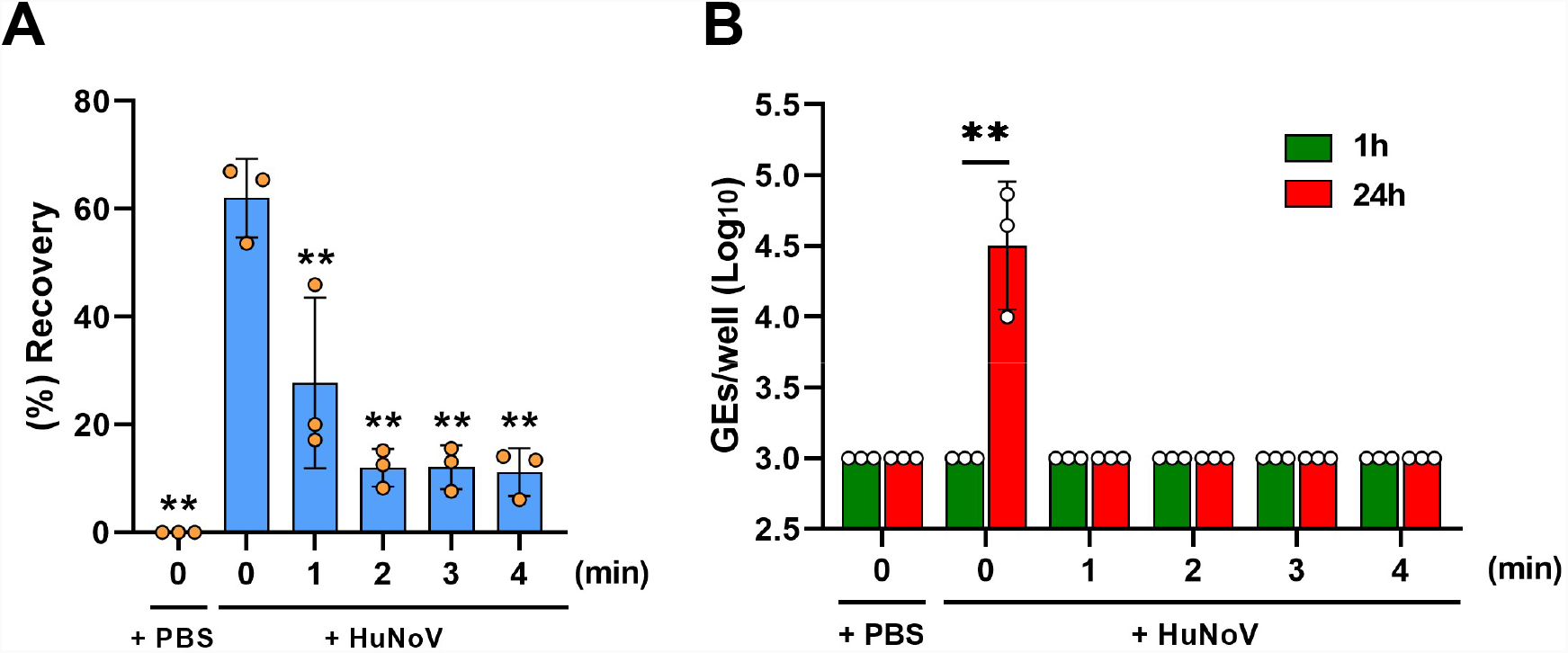
Thermal inactivation of clams artificially contaminated with HuNoV. The clam tissue was spiked with either PBS or HuNoV containing stool filtrate. The spiked clam tissue was next left-untreated or heat-treated at 90 °C for 1, 2, 3, or 4 minutes, and then homogenized. (A) Viral GEs in the clam extracts were quantified by RT-qPCR and the recovery efficiency was calculated as in Materials and methods. ** *p* < 0.01 *versus* unheated HuNoV-spiked clam samples (0 min), one-way ANOVA followed by Dunnett’s multiple-comparison test. (B) The samples were inoculated to differentiated HIEs and HuNoV GEs in the HIEs were determined as in Materials and methods. ** *p* < 0.01, two-tailed Student *t* test. Results were shown as mean ± standard deviation calculated from 3 independent experiments (n=3).

Then, infectious virus was quantified by determining viral GEs at 1 or 24 hpi as described in the Materials and methods section. HuNoV in clam homogenates was infectious and able to replicate in HIEs showing a 42.5-fold increase in GEs at 24 hpi as compared to 1 hpi, while PBS injected samples contained no infectious virus as expected (Fig. 2B). Furthermore, all groups of heat treatment showed no viral replication at 24 hpi, demonstrating that the 90 °C for 1 minute treatment is sufficient for inactivation of HuNoV infectivity in clam tissue homogenates (Fig. 2B).

## Discussion

Generally, if no cultivation system is available to grow a certain human pathogen, employing cultivable surrogate virus(es) is used to study the biological characteristics including tolerability against disinfectants or heating (Bozkurt et al., 2015). However, the properties of the surrogate viruses are not always the same as the human pathogen. Indeed, previous reports demonstrate that HuNoV is resistant to 70% alcohol (Costantini et al., 2018), whereas surrogate viruses such as murine norovirus (MNV), feline calicivirus (FCV) and porcine enteric calicivirus, but not Tulane virus, are sensitive to this treatment (Cromeans et al., 2014). A study on heat inactivation of MNV in shellfish showed that the heat treatment at 90 °C for 90 seconds resulted in an approximate 2 log10 reduction of infectious virus measured by using plaque assays (Sow et al., 2011). Whether the heat treatment at 90 °C for 1 minute is sufficient for the inactivation of MNV as well as HuNoV remains to be verified.

Recently, a propidium monoazide (PMA)-viability RT-qPCR assay, which is expected to measure only infectious virions containing intact viral RNA, but not non-infectious virions containing degraded RNA, was applied to study inactivation of HuNoV in clams (Fuentes et al., 2021). That study demonstrated that HuNoV spiked in clams and heated at 90 °C for 10 minutes resulted in a 3.52 log10 reduction of HuNoVs determined by PMA-viability RT-qPCR assay (Fuentes et al., 2021). Since growth of HuNoVs in HIEs is still lower than that of surrogate viruses in their cultivable cells, improvement of growth efficiency is required to be utilized as general evaluation methods for HuNoV inactivation. For example, conditions to cultivate HuNoV in genetically modified lines and with improved medium conditions to achieve at least 3 logs of replication are being sought for optimal inactivation studies (Lin et al., 2020; Ettayebi et al., 2021). Meanwhile, comparisons between the PMA-viability RT-qPCR assay and our evaluation method using HIEs would be beneficial to develop a general pragmatic method to evaluate HuNoV inactivation.

In this study, we used freshwater clams to evaluate HuNoV inactivation using HIEs, because the HuNoV genome is frequently detected in clams as well as oysters among bivalve mollusks (Guix et al., 2019). It would be worth comparing HuNoV inactivation patterns between in clams and oysters, which needs further investigation.

In summary, we evaluated the heat inactivation of HuNoV in freshwater clams using HIEs and showed that treatment at 90 °C for 1 minute is sufficient for inactivation of HuNoV in the clam bodies. We provided direct evidence regarding HuNoV inactivation in a contaminated food using the HIE culture system. This information will be valuable to develop guidelines to inactivate HuNoV, which will contribute to reducing the risk of foodborne illness associated with HuNoV.

## Data availability statement

The original contributions presented in the study are included in the article, further inquiries can be directed to the corresponding author.

## Author contributions

K.M. conceived and supervised the project. K.M. and T.H. designed the experiments and wrote the manuscript. T.H., Y.Y. and K.M. conducted the experiments and analyzed the results. A.T., T.K., M.M., and M.K.E. provided advice for the study, discussed the results, and critically reviewed the manuscript. All authors approved the final version of the manuscript.

## Acknowledgements

We thank the research group of freshwater crams in Japan for kind support for cram handling.

